# The dorsal hippocampus is required for the formation of long-term duration memories in rats

**DOI:** 10.1101/2020.12.13.421222

**Authors:** Toshimichi Hata, Tatsuya Yamashita, Taisuke Kamada

**Affiliations:** Faculty of Psychology, Doshisha University, 1-3 Tatara Miyakodani, Kyotanabe, Japan 610-0394; Graduate School of Psychology, Doshisha University, 1-3 Tatara Miyakodani, Kyotanabe, Japan 610-0394; Medical Innovation Center, Graduate School of Medicine, Kyoto University, Sakyo-ku, Kyoto, Japan 606-8507

**Keywords:** Interval timing, Time perception, Peak interval, Dorsal hippocampus, Long-term memory, Short-term memory, Time-shift paradigm

## Abstract

Interval timing—the perception of durations mainly in seconds or minutes—is a ubiquitous behavior in organisms. Animal studies have suggested that the hippocampus plays an essential role in duration memory; however, the memory processes involved are unclear. To clarify the role of the dorsal hippocampus in the acquisition of long-term duration memories, we adapted the “time-shift paradigm” to a peak-interval procedure. After a sufficient number of training with an initial target duration (20 s), the rats underwent “shift sessions” with a new target duration (40 s) under a muscimol (0.5 μg per side) infusion into the bilateral dorsal hippocampus. The memory of the new target duration was then tested in drug-free “probe sessions,” including trials in which no lever presses were reinforced. In the probe sessions, the mean response rate distribution of the muscimol group was located leftward to the control group, but these two response rate distributions were superimposed on the standardized time axis, suggesting a scalar property. In the session-by-session analysis, the mean peak time (an index of timing accuracy) of the muscimol group was lower than that of the control group in the probe sessions, but not in the shift sessions. These findings suggest that the dorsal hippocampus is required for the formation of long-term duration memories within the range of interval timing.

## INTRODUCTION

Interval timing—the perception, estimation, and discrimination of durations in seconds, minutes, or hours—is a fundamental ability in many species to perform behaviors such as foraging and decision making (Buhusi & Meck, 2005). For instance, the nectar-feeding birds Amakihi (*Loxops virens*) periodically revisit the flowers from which they had previously collected nectar for efficient foraging (Kamil, 1978). Pigeons and rats underestimate the value of delayed reinforcement as a function of the duration of the delay (Mazur, 1987; Richards, Mitchell, & de Wit, 1997; Green, Myerson, Holt, Slevin, & Estle, 2004). For humans, instant noodles would be soggy if we could not estimate the interval it takes for them to be ready. Thus, interval timing is a ubiquitous behavior in organisms.

Animal studies have shown that hippocampal dysfunction alters interval timing behavior (for review, Lee, Thavabalasingam, Alushaj, Cavdaroglu, & Ito, 2019). The “classic effects” (Yin & Troger, 2011) of hippocampal or hippocampus-related regions dysfunctions on interval timing are to alter the accuracy (the length of the estimated or “subjective” durations) in the peak-interval (PI) procedure (Hata & Okaichi, 1998; Meck, 1988; Meck, Church, & Olton, 1984; Olton Meck, & Church, 1987; Tam, Jennings, & Bonardi, 2015; Yin & Meck, 2014) and the differential reinforcement of low rate (DRL) schedules (Jaldow & Oakley, 1990). These findings suggest that hippocampal lesions alter the accuracy of interval timing, but do not impair the ability to regulate behaviors depending on the passage of time.

Other lines of evidence from trace conditioning, electrophysiology, and computational models suggest that the hippocampus and hippocampal neurons are involved in the memory or coding of durations (see Table 1 in Lee et al., 2019). In trace conditioning, an unconditioned stimulus (US) is delivered after the termination of a conditioned stimulus (CS). Many studies using CS offset - US onset intervals (CS-US intervals) in the seconds range (i.e., range of the interval timing) have reported that hippocampal lesions, especially on the dorsal parts, impair the acquisition and retention of trace fear conditioning (McEchron, Bouwmeester, Tseng, Weiss, & Disterhoft, 1998; McEchron, Tseng, & Disterhoft, 2000; Quinn, Oommen, Morrison, & Fanselow, 2002; Quinn, Loya, Ma, & Fanselow, 2005; Lin & Honey, 2011; Raybuck & Lattal, 2011; Sellami, Abed, Brayda-Bruno, Etchamendy, Valério, Oulé, … Marighetto, 2017). Lee et al. (2019) stated that trace conditioning studies largely substantiate the hippocampus in representations of temporal duration memory. The hippocampal “time cell,” which fires at a specific moment in a temporally structured experience, has received much attention in the last decade (Pastalkova, Itskov, Amarasingham, & Buzsaki, 2008; MacDonald, Carrow, Place, & Eichenbaum, 2013; Saltz, Tiganj, Khasnabish, Kohley, Sheehan, Howard, & Eichenbaum, 2016; for review, Eichenbaum, 2014). Moreover, recent computational studies have suggested that the temporal distance between the to-be-learned time criterion and the time of the peak activity of each time cell provides an error signal to correct the synaptic weight (Oprisan, Aft, Buhusi, & Buhusi, 2018), and the memory of the duration is stored in the hippocampus (Oprisan, Buhusi, & Buhusi, 2018). Taken together, these recent findings suggest that the hippocampus plays an important role in duration memory.

**Table 1.**
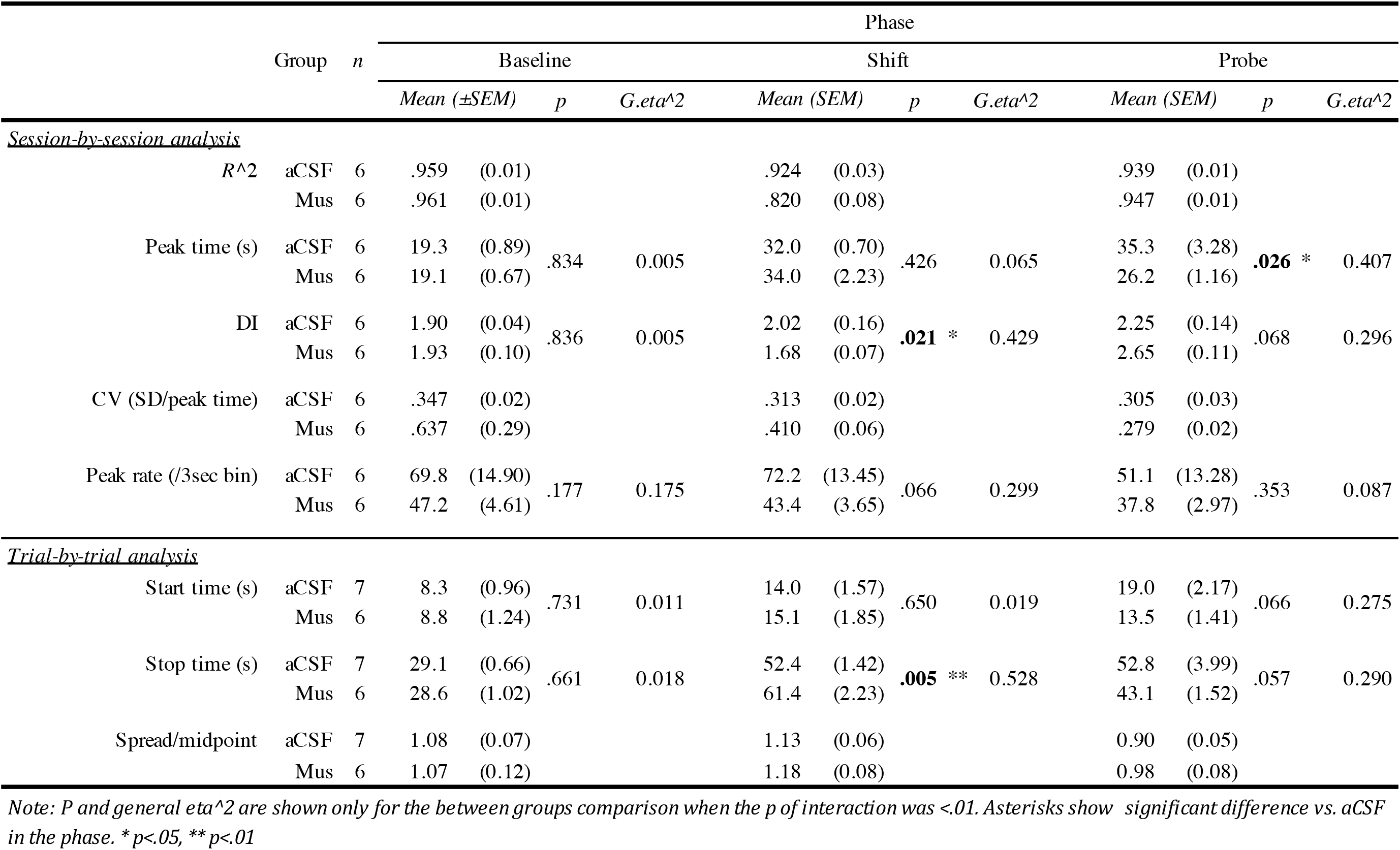
*Mean(± SEM)*, *p*, and *general eta^2* after excluding outliers.

However, these studies are insufficient to determine whether the hippocampus plays a vital role in the acquisition of duration memories. First, studies on trace conditioning did not selectively focus on duration memory. Thus, the impairment in the acquisition of trace conditioning following hippocampal dysfunction can be explained not only by the inability to acquire the memory of the duration but also by other impairments, such as the disappearance of the memory trace of the CS presentation, or impairment of the “continuation” of the timing proposed in the ICAT model of temporal processing (Petter, Lusk, Hesslow, & Meck, 2016). In the ICAT model, it is assumed that the interval timing is completed in four distinct phases: initiation, continuation, adjustment, and termination of timing. Continuation is defined as “monitor the passage of time during fill or unfilled intervals” (Petter et al., 2016). In trace conditioning, animals have to monitor the CS-US interval to perform the conditioning. The authors insisted that human patients with hippocampal lesions demonstrate a capability for initiation and termination of timing, but an impairment in continuation (Petter et al., 2016). Second, due to the nature of electrophysiological correlational studies, it cannot be concluded in principle that the activity of the time cell is the source of the duration memory. Lesion or inactivation studies should be performed to answer this question. Taken together, an experiment that improves these deficiencies should be designed to determine whether the hippocampus plays an important role in the acquisition of duration memories.

To meet these requirements, we applied the “time-shift paradigm” (Hata, 2011) to the peak interval (PI) procedure. In the time-shift paradigm, after sufficient training with the initial target duration (e.g., 20 s), the target duration was changed to a different one (e.g., 40 s) in a “shift session” during which animals form a new duration memory. Previous studies reported that intra-striatal injection of the protein synthesis inhibitor, anisomycin, impaired the formation of duration memory (MacDonald, Cheng, & Meck, 2012) and Arc, a plasticity-related protein level in the striatum, was altered in the shift session (Dallérac, Graupner, Knippenberg, Martinez, Tavares, Tallot, … Doyère, 2017). We believe that this paradigm can dissociate the effects of a drug with regard to the acquisition of duration memory from its impact on other processes (e.g., task rules, which may be included in the early stage of learning during the first target duration).

As an important modification, we prepared probe sessions including trials in which no lever press was reinforced on the day after the shift sessions. An early study applying the time-shift paradigm to the PI procedure did not report impairments in the acquisition of the second duration memory in hippocampal-lesioned rats (Meck, 1988). However, this finding does not necessarily mean that the hippocampus does not play a crucial role in acquiring duration memory. In general, the PI procedure is comprised of two types of trials— the “food” and “empty” trials (for detail, see Figure 2A). Food trials are a discrete fixed interval (FI) schedule that allow learning of the target durations. Empty trials are used to test the extent of duration memory. Only data from empty trials are usually analyzed. In empty trials, the subjects can access not only the long-term duration memories that are acquired through the previous sessions but also the short-term duration memories that are retrieved from the recent food trials within the session. In such situations, even if the hippocampal lesion impairs the formation of long-term duration memory, the animals might behave adaptively in the shift sessions, depending on the short-term duration memory, if any. On the contrary, if probe sessions including only empty trials are prepared after the shift sessions, then the within-session short-term memory is no longer unavailable in the sessions. All the available memories in the probe sessions are limited to the long-term memories acquired through the previous sessions. Here, if hippocampal dysfunction impairs the acquisition of long-term duration memory in the shift sessions, the effects should come to the surface in the probe sessions.

**Figure 2.**
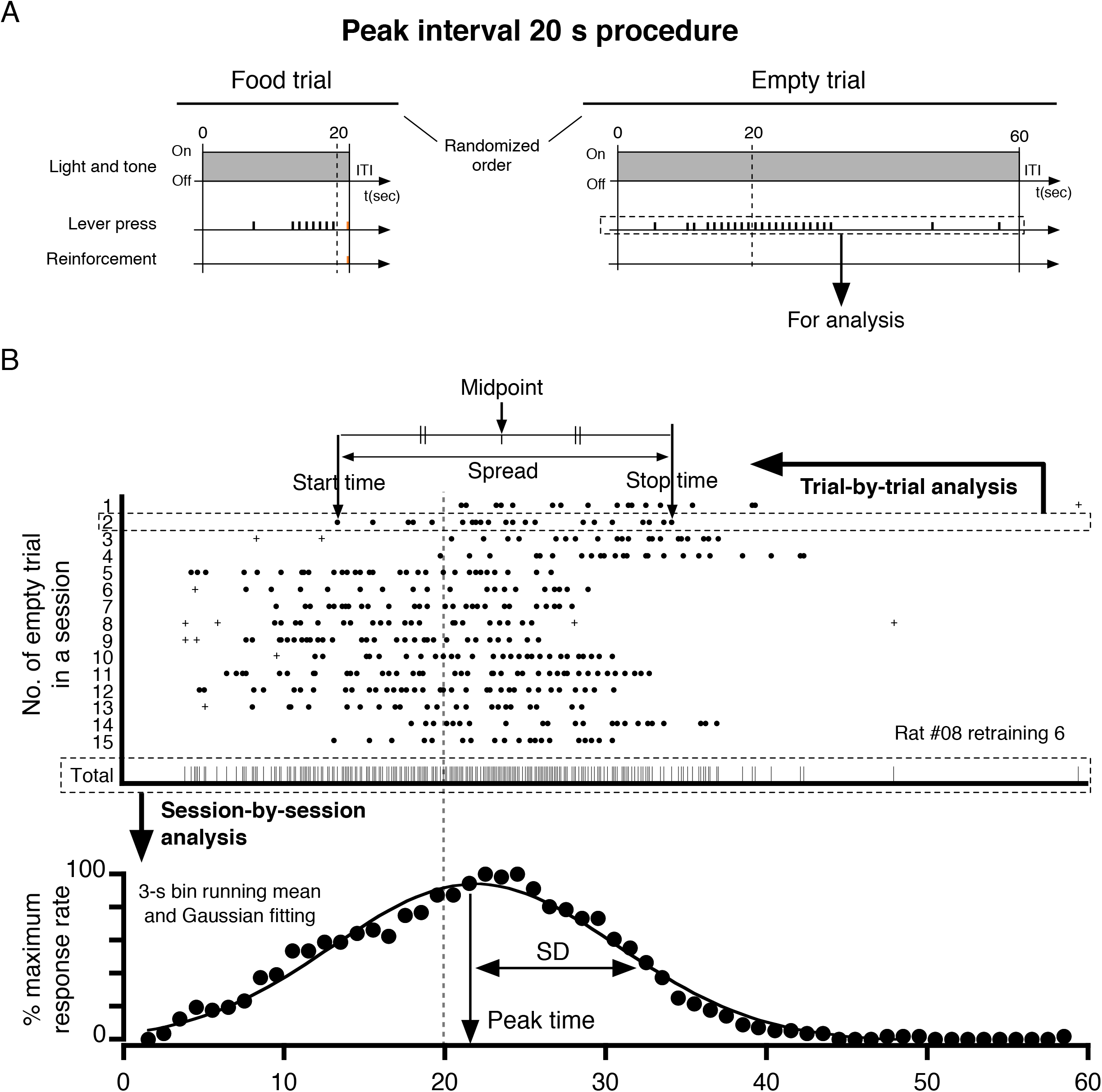
**(A)** Two types of trials included in the peak interval (PI) procedure. Only the data from empty trials were used for analysis. **(B)** A raster plot of lever-press responses (upper panel) and response rate distribution (lower panel) in a representative rat. In the trial-by-trial analysis, the start and stop times were identified using the algorism by Church et al. (1994). The crosses (+) show sporadic responses detected by the algorism.

Therefore, the purpose of the present study was to clarify the role of the dorsal hippocampus in the acquisition of long-term duration memory by applying the time-shift paradigm to the PI procedure with probe sessions. We hypothesized that an intra-hippocampal infusion of muscimol before the shift sessions would impair the acquisition of long-term memory of the second target duration; however, the effects would be detected in the probe sessions, but not in the shift sessions.

## METHODS

### Subjects

Sixteen experimentally naïve, 10-week-old male Wistar albino rats (Shimizu Laboratory Supplies, Kyoto, Japan) were used, but one animal died during surgery. Each rat was housed individually in a stainless-steel cage. The rats were deprived of food for the first day of shaping; hence, their body weight decreased to 85% of their free-feeding weight at the start of the behavioral procedure. To compensate for the natural growth, the body weight of the rats was increased at a rate of 5 g per week. Water was provided *ad libitum*. The temperature of the breeding room was maintained at 22 ± 2 °C. The light-dark cycle was 12:12 h, and the light phase started at 8:00 A.M. All experimental sessions were executed during the light phase. All experimental procedures were approved by the Doshisha Committee of Animal Experiments (A17075).

### Apparatus

Eight identical operant chambers were individually installed in a sound-attenuating box with a 25 W LED light on the ceiling. Each chamber (220 × 165 × 200 mm in DWH) was equipped with a non-retractable lever on the left side (40 mm from the floor) and a food cup (15 mm from the floor) in the center of the front wall. The floor was made of stainless steel bars with a 3 mm diameter, each bar 10 mm apart. A self-made application developed with XOJO^®^ (XOJO Inc. Austin, TX, USA) was run on two Macintosh PowerBook Air computers (Apple, Cupertino, CA, USA) to control the experiment and collect data. This was interfaced with two programmable controllers (SYSMAC CPM1A-40CDR-A-V1, OMRON, Kyoto, Japan) and two USB I/O controllers to control the eight chambers (RBIO-2U, Kyoritsu Electronic Industry, Osaka, Japan).

### Procedure

The flow of the experiments is shown in Figure 1.

**Figure 1.**
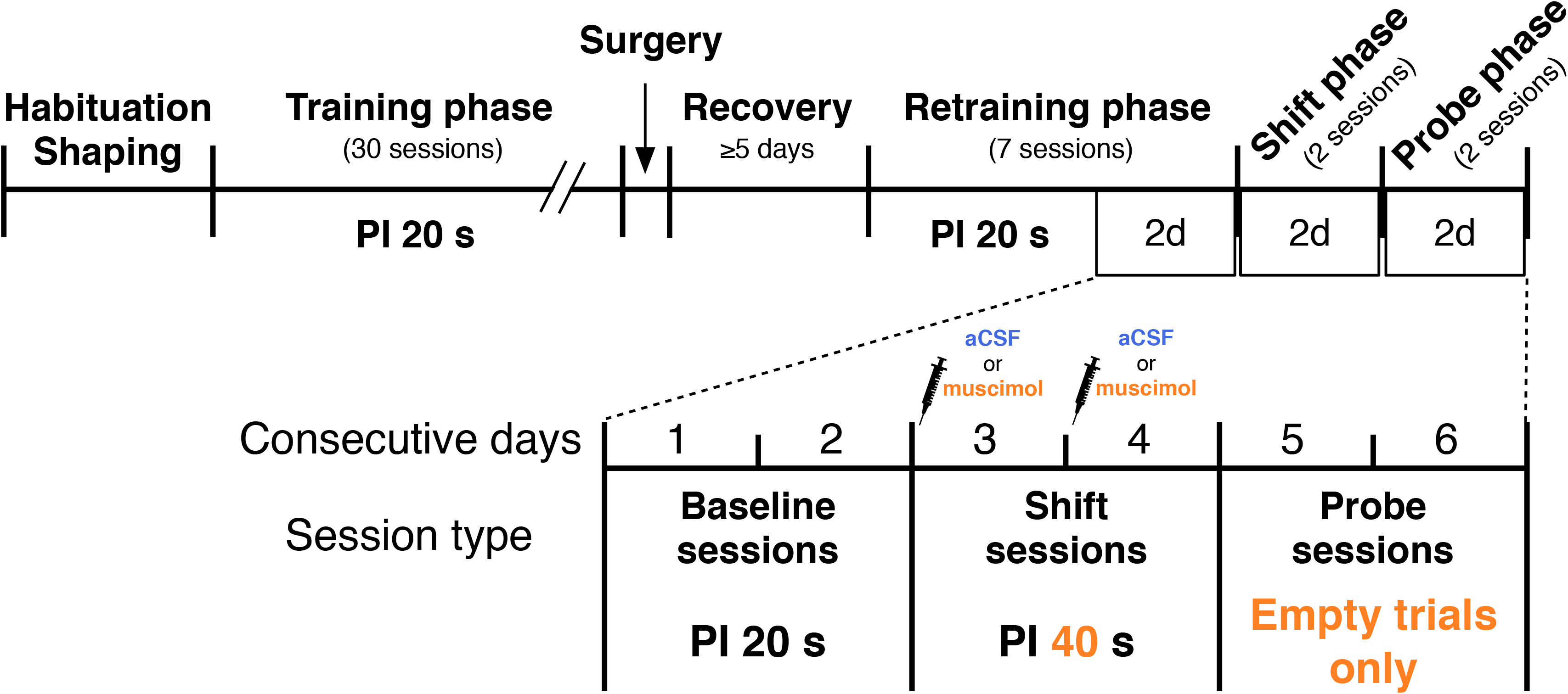
The flow of the experiment. The last two sessions of the retraining phase were defined as the baseline sessions. The baseline, shift, and probe sessions were executed over six consecutive days. Note that the required time was changed from 20 s to 40 s in the shift sessions, and that the drugs were infused before starting these sessions. In addition, only empty trials were included in the probe sessions.

#### Habituation

After handling for 5 min per day for 5 days, each animal was trained on a 20 s fixed time (FT) schedule for five sessions, one session per day. In each session, 30 food pellets (F0021-J, Bio-serv, Flemington, NJ, USA) were delivered separately every 20 s, irrespective of animal behavior. The house light was continuously turned on during the session.

#### Shaping

Rats were trained to press the lever under a continuous reinforcement schedule until they earned 60 pellets from their lever presses or after 60 min had passed in each session, whichever came first. The house light was turned on during the session. The acquisition criteria were that they earned 60 pellets within 60 min in three successive days.

#### PI 20 s training (sessions 1-30)

The rats were then trained using the PI procedure with one session per day (Figure 2A). A session consisted of two types of trials, food and empty trials, in a random order separated by intertrial intervals (ITIs) of 40 ± 10 s. The ratio of food to empty trials was 42:18 in sessions 1–21 and 15:15 in sessions 22–30. Both types of trials started with the onset of the house light and tone (signal stimuli) and finished with their offsets. In the food trial, the first lever press after 20 s from the start of the trial (orange ticks in Figure 2A) was reinforced with a food pellet, and then the trial was finished (discrete trial FI 20 s). An empty trial lasted for 60 s without any reinforcement for lever presses. If a lever press was emitted during the last 5 s of the ITI, the ITI was extended for 10 s from the press.

#### Surgery

Rats were anesthetized with 2–3% isoflurane (2-3 L/min flow rate, MK-A110, Muromachi, Tokyo, Japan) and fixed on a stereotaxic frame (Kopf Instruments, Tujunga, CA, USA). A guide cannula (C232G-3.8/SPC, Plastics One, Roanoke, VA, USA) attached to a dummy cannula (C232DC-3.8/SPC, Plastics One) was lowered into each hemisphere of the dorsal hippocampus. The target coordinates of the tip of the guide cannula based on a brain map (Paxinos & Watson, 1998) were as follows: −3.6 mm anteroposterior to the bregma, ± 1.9 mm mediolateral, and −2.2 mm dorsoventral from the skull surface. The cannula assembly was fixed to the skull surface using dental cement and two small screws. A dummy cannula was inserted into the guide cannula and fixed using a dust cap (303DC, Plastics One). Rats were allowed to recover for ≥5 days after the surgery. For a few days during the early recovery period, an antibiotic agent (Mycillinsol, Meiji, Tokyo, Japan) was applied to the wound once per day to avoid infection.

#### PI 20 s retraining (sessions 31-37)

After the recovery period, the rats were retrained using the PI procedure (seven sessions), similar to the training phase. The ratio of the food to empty trials was 15:15. The last two retraining sessions were defined as the baseline sessions (Figure 2A).

#### Shift session (sessions 38-39)

Two consecutive shift sessions were conducted from the day after the second baseline session. Rats were assigned to one of two groups, the artificial cerebrospinal fluid (aCSF) or muscimol (Mus) group, to ensure that the performances during the last three sessions of the retraining phase were as even as possible. Immediately before starting each session, the dummy cannula was replaced with an internal cannula (C232I-3.8/SPC, Plastics One), extending 1 mm from the tip of the guide cannula for infusion. The muscimol solution for Mus rats (0.5 μg/μL, 1 μL per side) or aCSF for aCSF rats (1 μL) was infused into the bilateral dorsal hippocampus at a flow rate of 0.5 μL/min for 2 min using a gastight syringe (100 μL, Hamilton, Reno, NV, USA) and microsyringe pump (ESP-32, Eicom, Kyoto, Japan). The cannula was left in place for 1 min to allow dispersion of the solution, which was then replaced with a dummy cannula. The dummy cannula was again inserted into the guide cannula and fixed with a dust cap. The significant changes in the procedure from the previous sessions were that lever press behavior was reinforced with a 40 s FI during food trials, and that the duration of the empty trials was extended to 120 s. The other parameters were the same as those used in the retraining sessions.

#### Probe session (sessions 40-41)

Two consecutive probe sessions were conducted on the day after the second shift session. The rats were tested in two sessions of 15 empty trials without infusions. The other parameters were the same as those used in the shift sessions.

### Dependent variables

#### Session-by-session analysis

For each session of each rat, the total number of lever-press responses in each 3-s bin (such as 0–3 s and 1–4 s) was counted over all empty trials (raw numbers of responses). To cancel the individual differences, they were converted to the percentage of the maximum number of responses among the bins (response rate). The response rates were plotted as a function of elapsed time (response rate distribution, Figure 2B, lower panel). Two consecutive response rate (as well as the raw response values) distributions within a phase (baseline, shift, and probe) were averaged. Only for the averaged response rate distributions the values were again converted to the percent maximum and plotted for fitting. These distributions were fitted with a Gaussian curve using the KaleidaGraph^®^ (ver. 4.5.3, Synergy Software, Reading, PA, USA): *R(t)* = *a* + b × exp { − .5 × [(*t* − *c*)/*d*]^2^}. Here, *t* is the time from the start of the empty trial, and *R*(*t*) is the response rate in bin *t*. Parameter *c* is the time at which the response rate was at its maximum. Thus, *c* was adopted as the peak time (an index of the subjective length of the to-be-timed duration), and d / √2 was defined as the standard deviation (*SD*). The CV (*SD*/*peak* time) was calculated as an index of timing precision (i.e., variability). The raw number of responses at the peak time was defined as the peak rate, an index of motivation (Roberts, 1981). The initial values of *a*, *b*, and *d* for the iterative estimation were 1, 100, and 10, respectively. To determine the initial *c*, the first and last bins, which had response rates of over 80%, were detected. The mean of the two class values of these bins was set as the initial *c*. The data used for fitting ranged from 1.5 s to 38.5 s for the baseline sessions and 1.5 s to 78.5 s for the shift and probe sessions, which were performed to cut off the noise observed in the last part of the empty trials. The allowable error of the interactive estimation was set to 0.1%. Curve fitting was also performed for the group mean response rate distribution. For the group mean distribution, the initial *a*, *b*, and *d* values were the same as for the individual distribution, whereas *c* was set at 20 and 40 for the baseline and the shift/probe sessions, respectively. The data used to fit the group mean distributions ranged from 1.5 s to 58.5 s for the baseline sessions and 1.5 s to 118.5 s for the shift and probe sessions. For the standardized mean response rate distributions (see Figure 4G-I), the values of the x (time) axis were converted to the ratio of the estimated peak time in each group. The original time values corresponding to the standardized x of 0.1 steps were substituted into the regression curve formula. The Y values were then converted to the maximum Y value.

**Figure 4.**
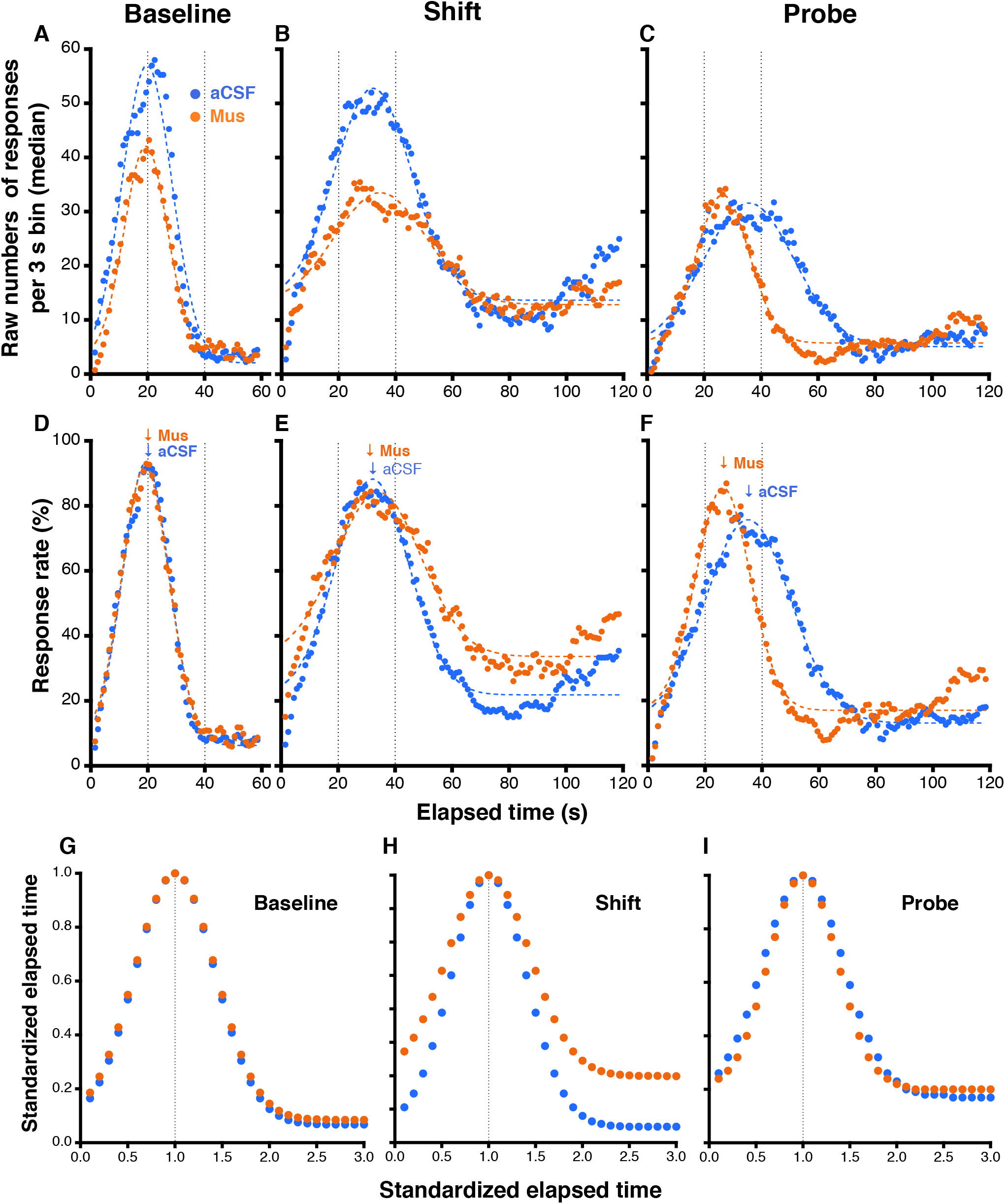
**(A-C)** The distribution of the median raw number of responses per 3 s bin in the baseline **(A)**, shift (B), and probe **(C)** sessions. Blue circles represent the aCSF, and orange circles represent the Mus group. The two vertical dotted lines indicate the required time of the baseline sessions (20 s) and the shift sessions (40 s). **(D-F)** Mean response rate distributions. The arrows show the time at which the response rate was at the maximum for each group. **(G-I)** Standardized mean response rate distributions. The horizontal axis was converted to the ratio to the peak time of each distribution. In the qualitative observation, the two curves overlapped as a whole in the baseline and probe sessions, but not in the shift sessions.

The discrimination index (*DI*) was also calculated for the individual response rate distribution for the fitting using *DI* = 100 / (mean response rate of all bins in the distribution) (Meck & Church, 1987) as an index of precision. For the *DI* calculation, the range of averaging in the denominator was from 1.5 s to 38.5 s for the baseline sessions and from 1.5 s. to 78.5 s for the shift and probe sessions.

#### Trial-by-trial analysis

In an individual empty trial, rats showed a low–high–low response pattern (Figure 2B, upper panel). That is, they typically started pressing the lever before the required time, kept responding, and then stopped responding after the required time. According to an algorithm proposed by a previous study (Church, Meck, & Gibbon, 1994), the timepoints of the transition from low to high (start time) and high to low (stop time) can be detected, ignoring sporadic responses (crosses in Figure 2B, upper panel). The start and stop times were the indices of timing accuracy, in parallel with the peak time of the session-by-session analysis. The midpoint was defined as the mean of the start and stop times. The spread calculated by stop time minus start time was divided by midpoint (spread/midpoint), which is an index of the timing precision, similar to CV in the session-by-session analysis.

### Histology

After completing all behavioral sessions, the rats were deeply anesthetized with sodium pentobarbital (130 mg/kg, i.p.) and infused with dye (2% pontamine sky blue) using a procedure similar to that of drug infusion to locate the tips of the internal canulae. They were then perfused intracardially with saline, followed by the ALTFiX^®^ (FALMA, Tokyo, Japan). Their brains were removed and soaked in 10% and 20% sucrose phosphate buffer (0.1 M) until they sunk in each solution. The brains were sectioned in the coronal plane (40-μm thickness) using a cryostat (CM1850, Leica, Wetzlar, Germany). The sections were stained with cresyl violet to assess the location of the cannula tip.

### Statistical analysis

Two-way mixed ANOVAs followed by the test of simple main effects and Holm’s multiple comparisons were performed using “anovakun” (ver. 4.8.5., http://riseki.php.xdomain.jp/index.php) and “ggplot2” (Wickham, 2016) in R software (ver. 4.0.0, R Core Team, Vienna, Austria). The significance level was set to α=.05.

## RESULTS

### Histological results

A photomicrograph of a representative section is shown in Figure 3A. The tips of the guide cannulae (arrows) were located above the dorsal hippocampus, while those of the internal cannulae were extended 1 mm from the guide cannula. Hence, all tips of the internal cannulae were presumed to be located within the dorsal hippocampus (Figure 3B, C). Apparent injuries beyond the cannula tracks were not observed. We could not find a systematic difference in tip location between the included and excluded animals (for additional analysis, see Figure 5C).

**Figure 3.**
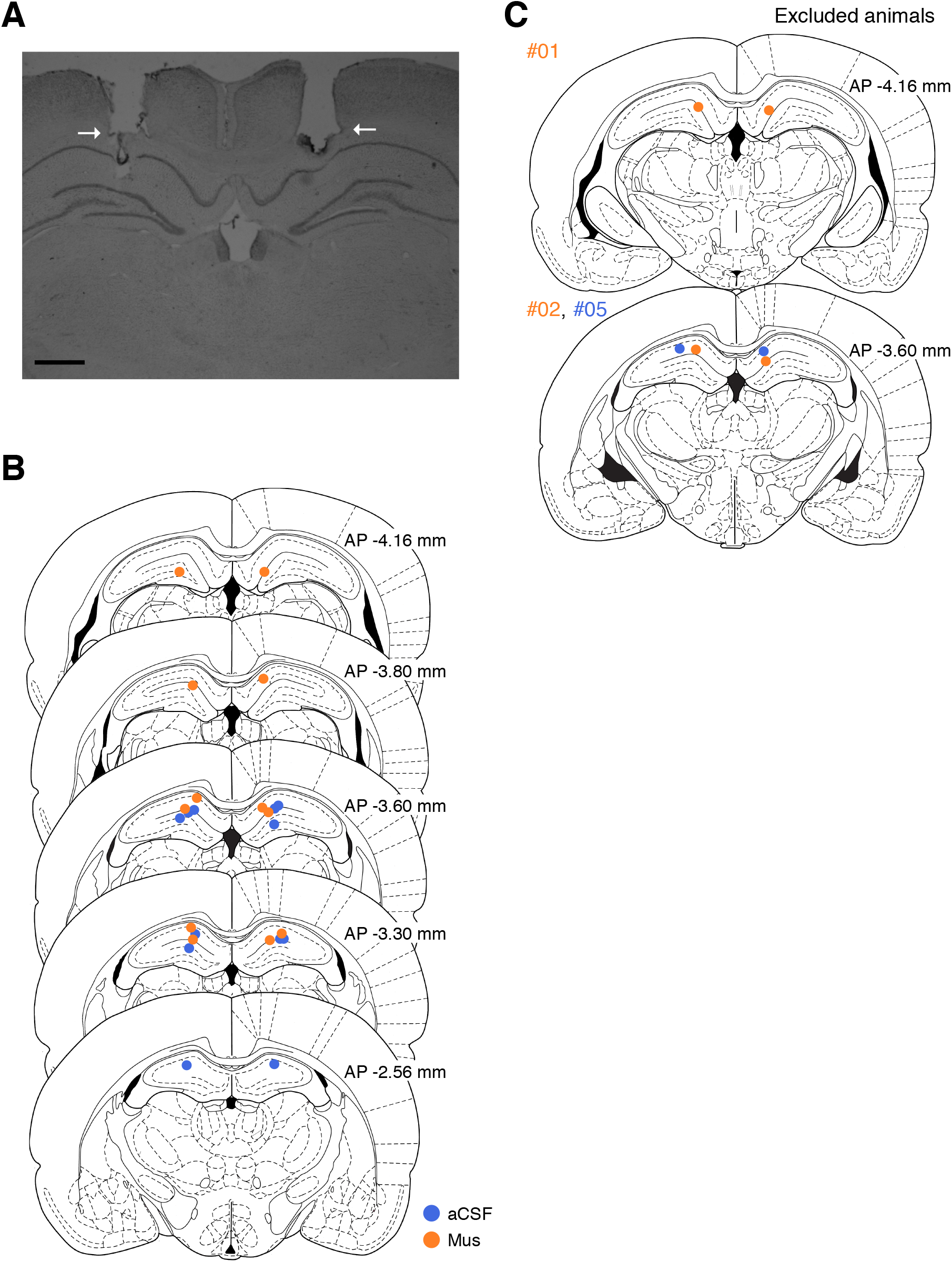
**(A)** A representative photomicrograph showing the tracks of the cannulae. The tips of the guide cannulae (arrows) were located above the dorsal hippocampus. The tips of the internal cannulae were extended 1 mm ventrally from these. Scale bar = 1 mm. **(B, C)** Placements of the internal cannula tips for the animals included in (B) and excluded from (C) data analysis. Blue circles represent the aCSF, and orange circles represent for the Mus rats. Reprinted from Paxinos, G. & Watson, C. *The Rat Brain in Stereotaxic Coordinates. 4th ed. [CD-ROM]*, Copyright (1998).

**Figure 5.**
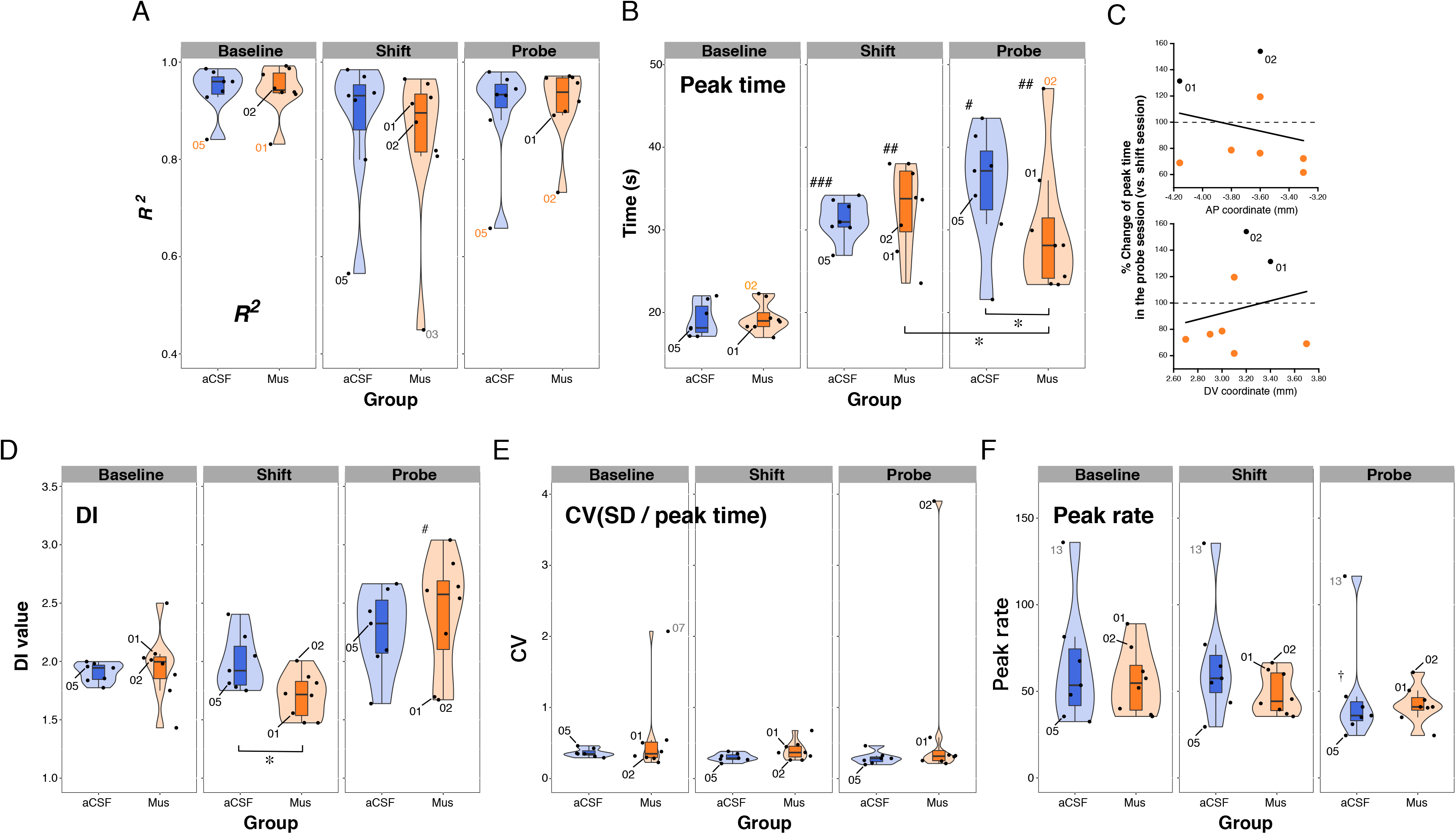
Violin plots of the *R*^*2*^ value of the fitting curve **(A)** and peak time **(B)**. Black-numbered points show the excluded animals. Red-numbered points indicate the data violating the criteria. Dimmed-numbered points are outliers but were included in the analysis as they did not violate the criteria. **(C)** Scatter plots of the percent change of peak time in the probe sessions to the shift sessions *vs*. AP coordinates (upper panel) and DV coordinates (lower panel) of the cannulae tips. Only the animal in the Mus group was plotted. Black circles represent excluded animals (01 and 02). Violin plots of DI **(D)**, CV (*SD* of fitted curve / peak time **(E)**), and peak rate **(F)**. ^*^*P* < .05. ^#^*P* <.05, ^##^*P* < .01, ^###^*P* < .001 *vs*. the baseline of the group. ^†^*P* < .05 *vs*. the baseline and the shift sessions.

### Session-by-session analysis

The data from three rats (01, 02, and 05) were excluded from the session-by-session analysis because the *R*^*2*^ or the peak time of the fitting curve of their baseline or probe sessions were over ± 2 *SD* from the mean of each group. As a result, the number of subjects used in the session-by-session analysis was six for each group.

#### Response rate distribution

To provide a qualitative overview of the performance, the distributions of the mean median of raw number of responses (Figure 4A-C), mean response rate (Figure 4D-F), and mean standardized response rate (Figure 4G-I) were plotted. In the raw number of responses, the height of the distributions in the aCSF group was higher than in the Mus group in the baseline and shift sessions, whereas they were similar to each other in the probe sessions. In the response rate, the distributions at baseline could be seen to overlap with each other (Figure 4D). The peaks in both groups were located at approximately 19 s. In the shift sessions (Figure 4E), although the Mus group had a wider distribution than the aCSF group, the peaks of both groups were located around 33 s. Interestingly, the distributions from the probe sessions (Figure 4F) showed a dramatic difference between the groups: The distribution from Mus group appeared to the left of that of the aCSF group. The peak of the Mus group was approximately 26 s, whereas that of the aCSF group was approximately 35 s. To evaluate the scalar properties, the standardized distributions are shown in Figure 4G-I. The distributions were superimposed as a whole in the probe sessions and at the baseline (Figure 4G, I), but not in the shift sessions (Figure 4H).

#### R^2^

To confirm the goodness of curve fitting, the group means of *R*^*2*^ (± *SEM*) were calculated (Table 1, distributions are shown in Figure 5A). All mean values were > .820. A mixed two-way ANOVA did not detect any significant effects (interaction and main effect of phase: *Fs*_1.05, 10.46_ = 2.00 and 4.42, *df*s were adjusted by Huynh-Feldt-Lecoutre’s epsilon, *Ps* = .187 and .059, *η*_*G*_^*2*^*s* = .094 and .187; main effect of group type: *F*_1, 10_ = .810, *P* = .389, *η*_*G*_^*2*^ = .038).

#### Peak time

To examine the effect on the accuracy of interval timing, group means (± *SEM*) of peak time were calculated (Table 1, distributions are shown in Figure 5B). There was no difference in the mean peak time of the baseline and shift sessions, whereas it was lower in the Mus group than in the aCSF group for the probe sessions. A mixed two-way ANOVA revealed that the interaction of group × phase (*F*_1.68, 16.82_ = 4.96, *P* = .025, *η*_*G*_^*2*^ = .267). The simple main effect of the group type was significant for the probe sessions (*F*_1, 10_ = 6.85, *P* = .026, *η*_*G*_^*2*^ = .407), but not for the baseline and shift sessions (*Fs*_1, 10_ = .05 and .69, *Ps* = .834 and .426, *η*_*G*_^*2*^*s* = .005 and .065, respectively). The simple main effects of the phase were significant in both groups (aCSF: *F*_1.03, 5.16_ = 15.55, *P* = .010, *η*_*G*_^*2*^ = .703. Mus: *F*_1.88, 9.41_ = 24.06, *P* < .001, *η*_*G*_^*2*^ = .767). The post-hoc Holm’s multiple comparisons test showed that the peak times of the shift and probe sessions were significantly higher than the baseline in each group. Moreover, in the Mus group, the mean peak time of the probe sessions was significantly lower than that of the shift sessions.

To confirm the relationship between cannula location and performance, scatter plots between the AP/DV axis of the cannulae and the percent changes of the peak time in the probe sessions to shift sessions are shown in Figure 5C. Pearson’s correlation coefficients were *r* = −.236 (*t*_*6*_ = −0.59, *P* = .574) and .209 (*t*_*6*_ = .52, *P* = .619) for AP and DV, respectively. Therefore, there was no significant relationship between the tip coordinates and the change in peak time.

#### DI

To examine the effect on the precision of interval timing, group means (± *SEM*) of DI were calculated (Table 1, distributions are shown in Figure 5D). There was no difference in *DI* between the baseline and probe sessions, whereas it was lower in the Mus group than in the aCSF group in the shift sessions. A mixed two-way ANOVA revealed a significant interaction (*F*_2, 20_ = 8.03, *P* = .003, *η*_*G*_^*2*^ = .27). The simple main effect of the group was significant in the shift sessions (*F*_1, 10_ = 7.52, *P* = .021, *η*_*G*_^*2*^*s* = .429), but not in the baseline and probe sessions (*Fs*_1, 10_ = .045 and 4.20, *Ps* = .836 and .068, *η*_*G*_^*2*^*s* = .005 and .296, respectively).

#### CV (SD/peak time)

As another index of the precision of interval timing, group means (± *SEM*) of CV were calculated as *SD/peak* time (Table 1, distributions are in Figure 5E). There were no noticeable differences between the groups in any of the phases. Mixed two-way ANOVA did not detect any significant effects (interaction and main effect phases: *Fs*_1.09, 10.85_ = 1.07 and 1.73, *Ps* = .331 and .217, *η*_*G*_^*2*^*s* = .053 and .084, respectively; main effect of group type: *F*_1, 10_ = 1.02, *P* = .336, *η*_*G*_^*2*^ = .046).

#### Peak rate

As an index of motivation, group means (± *SEM*) of the peak rate were calculated (Table 1, distributions are shown in Figure 5F). According to the distribution observations, an outlier seemed to increase the mean peak rate of the aCSF group. Mixed two-way ANOVA revealed a significant interaction (*F*_1.94, 19.4_ = 4.07, *P* = .034, *η*_*G*_^*2*^ = .019). The simple main effect of the phases was significant in the aCSF group (*F*_1.11, 5.55_ = 15.75, *P* = .008, *η*_*G*_^*2*^ = .085). Holm’s multiple comparisons test showed that the peak rate of the probe sessions was significantly lower than that of the baseline and shift sessions.

### Trial-by-trial analysis

Data from two rats (01 and 02) were excluded from the trial-by-trial analysis because the values of one of the three indices (start time, stop time, or spread) in the baseline or probe sessions were over ± 2 *SD* from the mean of each group. As a result, the numbers of subjects in the trial-by-trial analysis were 7 and 6 for aCSF and Mus, respectively.

#### Start time and stop time

As indices of timing accuracy, the group means of medians (± *SEM*) of the start times and the stop times over all trials in the phase (15 trials × 2 sessions, 30 trials) were calculated (Table 1, distributions are shown in Figure 6A, B). For the start time (Figure 6A), a mixed two-way ANOVA (group × phase) revealed that the interactions (*F*_1.60, 17.58_ = 3.215, *P* = .074, *η*_*G*_^*2*^ = .137) and the main effect of the group (*F*_1, 11_ = .692, *P* = .423, *η*_*G*_^*2*^ = .028) were not significant. The main effect of the phase was significant (*F*_1.60, 17.58_ = 3.215, *P* = .074, *η*_*G*_^*2*^ = .433). For the stop time (Figure 6B), the mean of the Mus group was higher than that of the aCSF group in the shift sessions, whereas the relationship was reversed in the probe sessions. A two-way ANOVA revealed that the interaction was significant (*F*_1.45, 15.94_ = 9.067, *P* = .004, *η*_*G*_^*2*^ = .355). The simple main effect of the group was significant in the shift session (*F*_1, 11_ = 12.292, *P* = .005, *η*_*G*_^*2*^ = .528), but not in the probe session (*F*_1, 11_ = 4.501, *P* = .057, *η*_*G*_^*2*^ = .290). The simple main effect of the phase was significant in both groups (aCSF: *F*_1.19, 7.15_ = 27.811, *P* < .001, *η*_*G*_^*2*^ = .769, Mus: *F*_1.29, 6.44_ = 125.949, *P* < .001, *η*_*G*_^*2*^ = .929). The post-hoc Holm’s multiple comparisons test showed that the stop time of the probe sessions was significantly lower than that of the shift sessions in the Mus group but not in the aCSF group.

**Figure 6.**
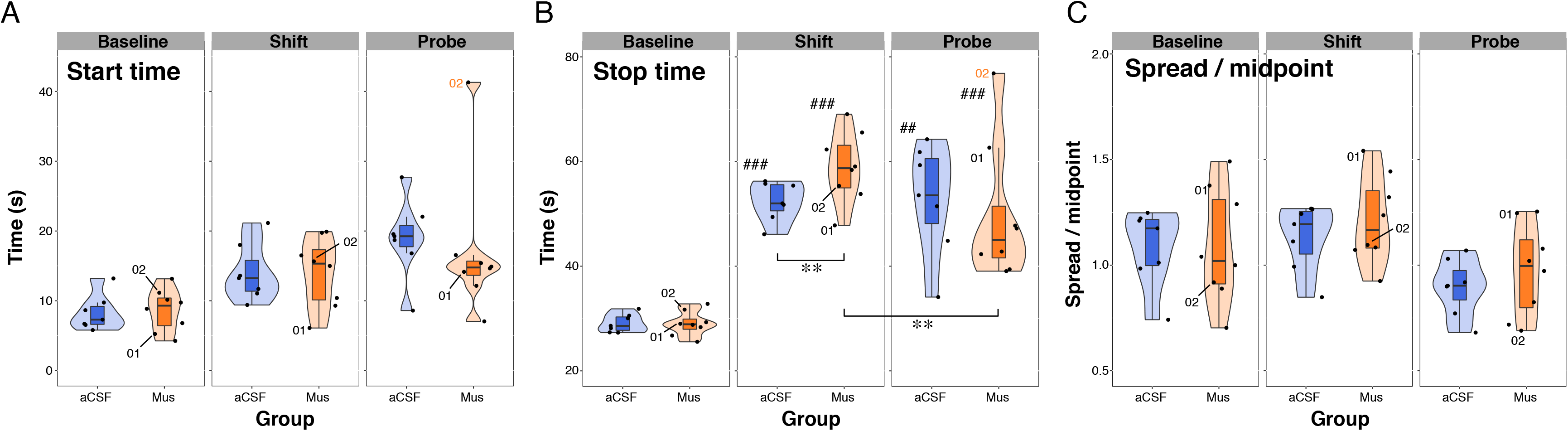
Violin plots of the start time **(A)**, the stop time **(B)**, and the spread/midpoint **(C)**. Black-numbered points show the excluded animals. Red-numbered points indicate the data violating the criteria. ^**^*P* < .01, ^##^*P* < .01, ^###^*P* < .001 *vs*. the baseline of the group.

#### Spread/midpoin

As an index of precision, group means (± *SEM*) of the spread/midpoint were calculated (Table 1, distributions are shown in Figure 6C). The interaction (*F*_2, 22_ = .526, *P* = .598, *η*_*G*_^*2*^ = .013) and the main effect of group (*F*_1, 11_ = .175, *P* = .684, *η*_*G*_^*2*^ = .011) were not significant, whereas the main effect of the phase was significant (*F*_2, 22_ = 10.50, *P* < .001, *η*_*G*_^*2*^ = .207). Holm’s multiple comparisons test showed that the spread/midpoint of the probe sessions was significantly lower than that of the shift sessions.

## DISCUSSION

The main results of our experiment can be summarized as follows: In the Mus group, the peak and stop times of the probe sessions were significantly decreased from the shift session. Moreover, the peak and stop times in the Mus group were significantly lower than those in the aCSF group during the probe sessions. These results indicate that the accuracy indices were influenced by muscimol. However, the effects on the indices of precision (the DI and the *SD/peak* time in the session-by-session analysis and the spread/midpoint in the trial-by-trial analysis) or motivation (the peak rate) were not detectable, at least in the probe sessions.

Two fundamental prerequisites, the homogeneity of the two groups before the important experimental operations and the validity of the task, were suggested to be guaranteed by the data. First, the response rate distributions of both groups almost entirely overlapped with each other in the baseline sessions (Figure 4D, G). Moreover, the mean *R*^*2*^, peak time, and *DI* in the session-by-session analysis (Figure 5), and the mean start time, stop time, and spread/midpoint in the trial-by-trial analysis (Figure 6) in the baseline sessions were all similar between the groups. These findings suggest that the differences between groups in the following analysis do not stem from the bias of the assignment of the subjects to the groups. Second, the task’s validity was confirmed by the significant transition of the mean peak time from the baseline sessions to the shift sessions (Figure 5B). In the aCSF group, the peak time of the shift session was significantly higher than that of the baseline, suggesting that the memory of the new target time was acquired by the animals. In addition, the extended peak time was maintained in the probe sessions, as evidenced by the non-significant difference between these two phases. These findings suggest that the memory of the new target duration was maintained for at least 24 h.

In the shift session, dysfunction of the dorsal hippocampus was suggested to impair the precision, but not the accuracy, of the interval timing. The significant extension of the stop time (Figure 6B) inevitably induced elevated response rates after the peak time in the response rate distribution of the Mus group (Figure 4E). This elevation could account for the significantly lower *DI* value seen in the Mus group (Figure 5D), suggesting impaired precision (discussed further below). On the contrary, hippocampal dysfunction did not seem to impair the accuracy of the interval timing at the shift sessions, as evidenced by the lack of a difference in peak time in the shift sessions (Figure 5B). This finding, however, does not contradict the “classic effect” (Yin & Troger, 2011, see the Introduction section) of the hippocampal lesion on the interval timing. Many studies have repeatedly reported that dysfunction of the hippocampus results in shortening of the peak time (Hata & Okaichi, 1998; Meck, 1988; Meck et al., 1984; Olton et al., 1987; Tam et al., 2015; Yin & Meck, 2014). In these studies, the effect has been reported in sessions in which the required time of the PI procedure was constant and appeared after chronic dysfunction of the hippocampus. However, in our study, the required time was changed, and the hippocampal function was reversibly inhibited in only two sessions. In this way, the situation in our study was different from those suggestive of “classic” hippocampal dysfunction effects. Therefore, our findings do not contradict those of previous studies.

The noticeable findings observed in the probe sessions can be explained by the notion that the dorsal hippocampus is involved in the formation of long-term, but not short-term, duration memories. The most important asset of the probe sessions was that the sessions included only empty trials. Thus, within-session short-term memory was not available for feedback control of the behavior in empty trials. All of the memories that were available to guide their behavior were limited to those acquired during the training/retraining sessions and/or the shift sessions. It is known that when the required time is increased during peak interval procedures, the peak times gradually increase in normal rats (Lejeune, Ferrarra, Simons, & Wearden, 1997; Meck, Komeily-Zadeh, & Church, 1984). These findings suggest that the shorter the peak time, the weaker the memory of the new required time. Our data were as follows: the response rate distribution of the Mus group was located to the left of the aCSF group (Figure 4C, F). The mean peak time of the Mus group was significantly lower than that of the aCSF group in the probe sessions (Figure 5B). In the Mus group, the peak time in the probe sessions was significantly lower than that in the shift sessions, while there was no significant difference in the aCSF group (Figure 5B). In the trial-by-trial analysis, the mean start and stop times of probe sessions in the Mus group were lower (although not significantly) than those in the aCSF group (Figure 6A, B); however, the *P* values were only slightly above the significance level (Table 1, *P* = .066 and .057, respectively). Moreover, the stop time of the Mus group significantly decreased from the shift sessions to the probe sessions (Figure 6B). These results cannot be explained by a motivational factor, because there was no significant difference in the peak rate, an index of motivation, between groups in all phases (Table 1 and Figure 5F). Taken together, these results consistently suggest that the animals in the Mus group had a poorer memory of the new required time (40 s) than the aCSF animals in the probe sessions. Other important results were that the mean peak times were similar between the groups in the shift sessions (Figure 5B). This can be interpreted to show that the within-session short-term memory from the food trials was available even though the muscimol was infused, which guided the animals’ adaptive behavior. Some previous studies support this “short-term/long-term dissociation hypothesis” (Haettig, Stefanko, Multani, Figueroa, McQuown, Wood, 2011; Lee & Kesner, 2003; Nagahara & McGaugh, 1992). In a study on the DNMTS task in the eight-arm radial maze, an intra-dorsal hippocampal infusion of muscimol impaired their intermediate-term (5 min) memory, but not their short-term (10 s) memory (Lee & Kesner, 2003). Intra-dorsal hippocampal muscimol infusion immediately after memory acquisition resulted in impairments in the performance of an object memory test conducted in mice 24 h after acquisition (Haettig et al., 2011). Moreover, a muscimol infusion before the acquisition of inhibitory avoidance into the septum, which has strong reciprocal connections with the hippocampus, disrupted retrieval after 48 h, but not 15 s. After 15 min of memory acquisition, the performance was halfway between the 48 h and 15 s conditions (Nagahara & McGaugh, 1992). Although the task used and/or the target region in our study were different from those of the reference studies, it is not surprising that the within-session short-term memory was available in our experiment. Taken together, our evidence strongly suggests that dysfunction of the dorsal hippocampus impairs the formation of long-term memory of a new duration.

The short-term/long-term dissociation hypothesis can also explain the discrepancy between our findings, the findings of a previous study, and the lower *DI* of the Mus group in the shift sessions. First, as mentioned in the Introduction, Meck (1988) reported that hippocampectomized rats did not show a delay in the acquisition of memories of the new target duration. In this experiment, however, the effect of the hippocampal lesion was examined in the shift sessions, each of which comprised both food and empty trials. If this hypothesis is correct, it is reasonable that the previous study did not report impairments in the acquisition of the new target memory. Therefore, this “discrepancy” may be ostensible. Second, the lower *DI* of the Mus group in the shift sessions might be explained by the incomplete short-term memory and the resulting dysfunction in feedback control. A previous study suggested that normal animals modulate when to start responding and when to stop in PI procedures, which were regulated according to the information from their own performances in recent food trials (Meck, 1988). The study also reported that this feedback control was impaired by the hippocampal lesion. Even if the within-session short-term memory seemed to be available in the Mus group, the availability might be lower than that of the aCSF group because, according to previous studies (Lee & Kesner, 2003; Nagahara & McGaugh, 1992), the inferred time window in which short-term memory was maintained may vary from several dozen seconds to several minutes. However, in our PI procedure, the animals must maintain a memory for at least 160 s (40 s for the mean duration of ITI + 120 s for the duration of an empty trial) for feedback control. Indeed, the contents of the memories are different from each other (durations, spatial information, and fear emotions); therefore, the estimation may be too simplified. However, it seems reasonable to infer that a weakness of the feedback control resulting from the available but weak short-term memory produced a fluctuation of the “low-high-low” pattern and then lowered the *DI* in the shift sessions. This speculation should be confirmed in future studies.

The standardized response rate distributions, the CV (*SD/peak* time), and the spread/midpoint in the probe sessions consistently suggest that the Mus rat normally performs interval timing. In the literature on interval timing, it has been repeatedly confirmed that two response rate distributions produced under different required times are superimposed on the standardized time axis relative to the peak time, that is, the scalar property (Church et al., 1994; Gibbon, 1977; Gibbon & Church, 1990, 1992; Malapani & Fairhurst, 2002). This means that the ratio of the width of the response rate distribution to the peak time is a constant (i.e., Weber’s law); i.e., the CV (*SD/peak* time) and the spread/midpoint are constant. In our data, the two standardized response rate distributions were superimposed on each other during the probe sessions (Figure 4I). Moreover, the CV (*SD/peak* time, Figure 5E) and spread/midpoint (Figure 6C) did not significantly differ between the groups in the probe sessions. This evidence suggests that the Mus rats had a normal interval timing based on their own target times, which were different from those of the aCSF rats.

The most noticeable progress of our study was its extension of previous knowledge from trace conditioning, electrophysiology, and computational models. First, data from trace conditioning were believed to be evidence of the role of the hippocampus in temporal duration memory (Lee et al., 2019). However, other explanations, such as the disappearance of the memory trace of the CS presentation or an impairment of the “continuation” proposed by the ICAT model of timing (Petter et al., 2016), can also explain the data. Using a timing-specific task (peak interval task) and a filled duration (i.e., the duration in which the to-be-timed stimulus is continuously presented) and confirming the normal interval timing in the probe sessions in the Mus group, we could exclude alternative interpretations of the data from the trace conditioning. Second, muscimol inactivation demonstrated a causal, but not correlational, relationship between the hippocampus and the acquisition of long-term duration memory. Third, our data provided experimental support to the computational models arguing the memory role of the hippocampus in interval timing (Oprisan, Aft, Buhusi, & Buhusi, 2018; Oprisan, Buhusi, & Buhusi, 2018).

In summary, the present findings strongly suggest that the dorsal hippocampus plays an important role in the formation of long-term duration memory in the supra-second range.

## Acknowledgments

We would like to thank Editage (https://www.editage.com) for English language editing.

## AUTHOR CONTRIBUTIONS

T.Y. conceptualized the study. T.H. and T.K. supervised the study. T.Y., T.K., and T.H. performed the experiments. T.H. and T.K. analyzed the data. T.H. and T.K. wrote the manuscript.

## DATA AVAILABILITY STATEMENT

The datasets are available from the open science framework (OSF). https://osf.io/fqngb/?view_only=6d21bea2cec74fa784a9cd79f3fe3774

